# Whole genome SNP typing to investigate methicillin-resistant *Staphylococcus* aureus carriage in a health-care provider as the source of multiple surgical site infections

**DOI:** 10.1101/047597

**Authors:** Chandler C Roe, Kimberly S Horn, Elizabeth M Driebe, Jolene Bowers, Joel A Terriquez, Paul Keim, David M Engelthaler

## Abstract

**Background:** Prevention of nosocomial transmission of infections is a central responsibility in the healthcare environment, and accurate identification of transmission events presents the first challenge. Phylogenetic analysis based on whole genome sequencing provides a high-resolution approach for accurately relating isolates to one another, allowing precise identification or exclusion of transmission events and sources for nearly all cases. We sequenced 24 methicillin-resistant *Staphylococcus aureus* (MRSA) genomes to retrospectively investigate a suspected point source of three surgical site infections (SSIs) that occurred over a one-year period. The source of transmission was believed to be a surgical team member colonized with MRSA, involved in all surgeries preceding the SSI cases, who was subsequently decolonized. Genetic relatedness among isolates was determined using whole genome single nucleotide polymorphism (SNP) data.

**Results:** Whole genome SNP typing (WGST) revealed 283 informative SNPs between the surgical team member’s isolate and the closest SSI isolate. The second isolate was 286 and the third was thousands of SNPs different, indicating the nasal carriage strain from the surgical team member was not the source of the SSIs. Given the mutation rates estimated for *S. aureus*, none of the SSI isolates share a common ancestor within the past 14 years, further discounting any common point source for these infections. The decolonization procedures and resources spent on the point source infection control could have been prevented if WGST was performed at the time of the suspected transmission, instead of retrospectively.

**Conclusions:** Whole genome sequence analysis is an ideal method to exclude isolates involved in transmission events and nosocomial outbreaks, and coupling this method with epidemiological data can determine if a transmission event occurred. These methods promise to direct infection control resources more appropriately.

## Background

Surgical site infections (SSIs) account for 31% of all healthcare-associated infections in the United States [1]. A critical risk factor for the development of a SSI is the severity of wound contamination during surgery [2, 3]. As one of the most predominant organisms colonizing the skin, *Staphylococcus aureus* is a common causative agent of SSI [4], and methicillin-resistant *S. aureus* (MRSA) accounts for 15% of reported SSIs [5]. The most significantly associated independent circumstance for the development of a *S. aureus* SSI is nasal carriage by the patient [3]. Studies that examined perioperative decolonization of patients using intranasal mupirocin showed a significant decrease in the risk of developing a SSI [6–8]. The risk of a *S. aureus* SSI associated with *S. aureus* nasal carriage or colonization of surgical team members is unknown.

The implementation of whole genome sequencing (WGS) in the clinical setting to closely analyze isolates has allowed for precise identification and characterization of putative transmission events among patients. WGS also allows for high resolution analysis of source tracing in hospitals which can have the potential to impact hospital-based outbreak investigations and infection prevention practices [9–12]. To our knowledge, however, no study has investigated MRSA carriage among surgical staff members as a source of MRSA transmission to surgical patients using WGST.

In this study, a hospital infection control team identified a cluster of MRSA SSIs from a single surgical team. Antibiotic susceptibility profiles suggested that the cases were related. The surgical team in question was screened for nasal carriage by the infection control team and one member tested positive for MRSA by PCR, as well as by culture. The surgical team member was decolonized per hospital protocol, and the MRSA decolonization was documented as successful, based on further nasal screening. In this study, we retrospectively examine the MRSA strain from the colonized surgical team member along with patient and control strains to determine whether it was the source of the cluster of SSIs using whole genome single nucleotide polymorphism (SNP) typing (WGST) [13–15]. These results demonstrate the utility of WGST for hospital molecular epidemiology and provide confirmation that the source of infection for three patients was not related to the surgical team member isolate.

## Methods

### Surveillance

From February 2010 to August 2011, 24 MRSA isolates were obtained from 18 patients and one surgical team member in a rural 280-bed hospital (Table 1). Isolates were obtained from leftover, in-house testing as approved by the Institutional Review Board of the participating institution. No patient consent was needed as samples obtained were isolates only. The isolates fell into three groups; the first group comprised the surgical team member’s isolate (suspected point source) and three patients’ SSI isolates. The three isolates were from patients who developed MRSA surgical site infections following surgical procedures by the same medical team. The surgical team in question was screened for nasal carriage and one member tested positive for MRSA by the GeneXpert Infinity MRSA PCR (Cepheid), and subsequently MRSA was cultured from the original swab providing the surgical team member isolate. The surgical team member’s isolate was tested and confirmed positive for *mecA*, an assay included in the GeneXpert PCR assay.

**Table 1.** **Clinical histories of patients with MRSA infections, including suspected index case**. Multiple samples taken from individuals are labeled as “a” and “b.”

| Patient | Isolates | Source | Year Collected |
| --- | --- | --- | --- |
| Surgical Team Member | 1 | Nasal | February 2011 |
| Case 2 | 1 | Blood | February 2011 |
| Case 3a | 3 | Abdominal fluid | September 2010 |
| Case 3b | - | Anal fistula | August 2011 |
| Case 3c | - | Sputum | August 2011 |
| Case 4 | 1 | Deep wound | July 2010 |
| Hospital Control 1 | 1 | Tissue | March 2010 |
| Hospital Control 2 | 1 | Incision | December 2010 |
| Hospital Control 3 | 1 | Joint fluid | September 2010 |
| Hospital Control 4a | 2 | Tissue | March 2010 |
| Hospital Control 4b | - | Deep wound | March 2010 |
| Hospital Control 5 | 1 | Tissue | November 2010 |
| Hospital Control 6 | 1 | Tissue | October 2010 |
| Hospital Control 7 | 1 | Deep wound | December 2010 |
| Hospital Control 8a | 2 | Abdominal fluid | February 2011 |
| Hospital Control 8b | - | Abdominal fluid | February 2011 |
| Healthcare Control 9a | 2 | Head | February 2011 |
| Healthcare Control 9b | - | Head | February 2011 |
| Healthcare Control 10 | 1 | Right stump | March 2011 |
| Healthcare Control 11 | 1 | Urine | February 2011 |
| Community Control 12 | 1 | Blood | March 2011 |
| Community Control 13 | 1 | Boil | October 2010 |
| Community Control 14 | 1 | Abscess groin | February 2011 |
| Community Control 15 | 1 | Excision site abdominal | March 2011 |

The second group consisted of 10 background isolates considered to have no association with the first group of samples but were also from SSIs, collected randomly from the regular sample flow at the hospital. The third group comprised 10 background samples, also randomly collected, from outpatients and patients admitted to the hospital with MRSA infections. These isolates were sourced from blood, sputum, wound and urine specimens. In all three groups of isolates, multiple samples from individual patients, labeled as “a” and “b”, were taken over time to serve as a control threshold for genetic similarity and relatedness.

### DNA sequencing

Isolates were grown on trypticase soy agar media and DNA was prepared using DNeasy Blood & Tissue Kit as described by the manufacturer (Qiagen, Valencia, CA) with the addition of lysostaphin to the gram-positive extraction protocol from Qiagen. The DNA samples were prepared for multiplexed, paired-end sequencing with a 500 base pair insert using Library Preparation Kit with Standard PCR Library Amplification (KAPA Biosystems, Woburn, MA). A 100 bp read paired-end run was used for all twenty-four isolates on the GAIIx sequencing platform (Illumina, Inc. San Diego, CA).

### SNP Phylogenetic Analysis

A phylogeny of all the MRSA isolates was constructed based on a matrix that tabulates all SNPs among the genomes of the isolates by their locus within a reference genome. The matrix was generated using the Northern Arizona SNP Pipeline (NASP) (http://github.com/TGenNorth/NASP/releases/tag/v0.9.6). NASP aligns DNA sequencing reads to a reference, identifies SNPs and filters SNP loci based on user-defined parameters as previously described[14–18]. NASP generated a matrix for this study by aligning sequence reads from the 24 isolates to the reference genome, FPR3757 (NCBI accession number: NC_007793), using Novoalign (Novocraft.com). Reads that mapped to multiple locations within the reference genome were excluded from the alignments as were reads containing insertions or deletions. SNPs were identified in the alignments using the Genome Analysis Toolkit (GATK) [19]. Only SNP loci found throughout all samples were used for the phylogenetic analysis. Additionally, SNPs had to be in >90% of the reads and have minimum 10X coverage to be included in a final matrix. MUMmer version 3.22 [20] was used to identify duplicated regions within FPR3757 and SNP loci within these regions were removed from the final analysis. The phylogenetic analysis of the high-quality core genome SNP matrix was performed with *MEGA* version 5.05 [21] using the maximum parsimony algorithm and bootstrap analysis with 1,000 replicates. Read depth statistics and reference coverage were determined from NASP. The whole genome sequence read files were deposited in the NCBI Sequence Read Archive under BioProject ID PRJNA255788. Read data were also used to identify the multi-sequence locus type (MLST) using SRST2 [22].

## Results

### Genomic Investigation

The sequence analysis provided appropriate depth and breath of sequence coverage across all isolates to conduct robust WGST analysis. Mean sequence read coverage of the reference genome FPR3757 for all samples was ≥95% with a mean read-depth of 103X. The phylogeny constructed was based on 18,520 parsimony informative SNPs and revealed two distinct clades (Figure 1). MLST analysis using WGS data identified these clades as two distinct sequence types: ST5, which is known to include a major clone (commonly referred to as PFGE type USA100) that is historically been associated with healthcare-acquired infections; and ST8, a major clonal group of largely community-acquired strains (containing the virulent clone that is commonly referred to as PFGE type USA300). ST5 and ST8 represent two distantly related lineages of *S. aureus* that are common in hospitals across the country [23, 24].

**Figure 1.**
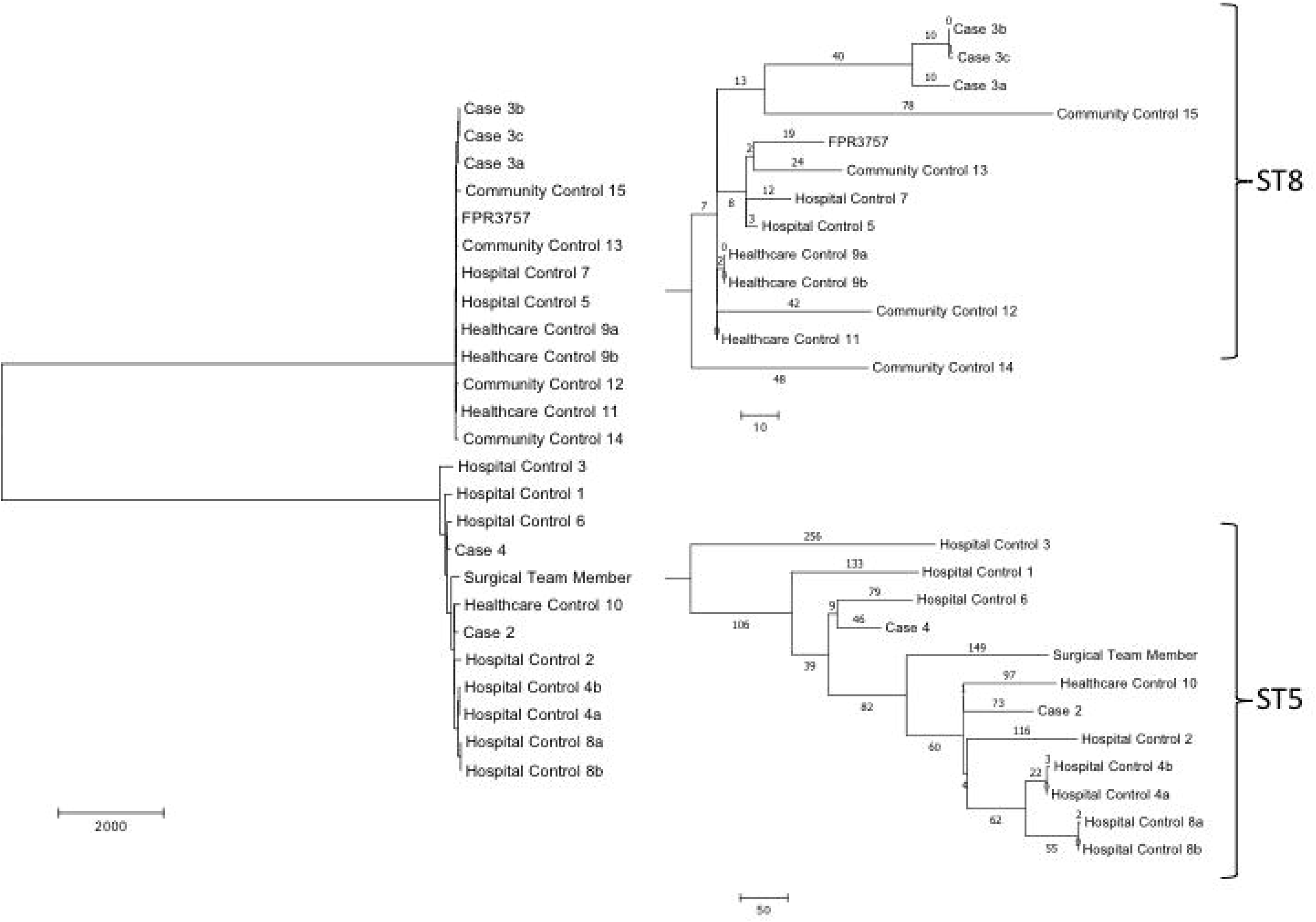
**Phylogenetic SNP analysis**. Unrooted phylogenetic SNP analysis based on whole genome sequence data of 24 MRSA isolates using the maximum parsimony algorithm.

WGST analysis demonstrated that no obvious transmission event occurred between the colonized surgical team member and patients. First, the two distinct clades in the phylogenetic tree, ST5 and ST8, are separated by 17,042 SNPs. The surgical team member isolate and one of the MRSA SSI cluster isolates, Case 3, are located in ST5 and ST8 respectively. Second, the surgical team member isolate is 283 SNPs different from the closest case isolate, Case 2, and even more distant from the third, Case 4. Estimates of the mutation rate for *S. aureus* have been calculated repeatedly at a rate of approximately 3.3 × 10^-6^ mutations per site per year, or one core genome SNP every 6 to 7 weeks [25–27]. Using this estimate, there should be approximately 8 to 10 SNPs per year after strains diverge from a common ancestor due to baseline mutation alone. Given their 283 SNP separation, the isolates from the surgical team member and Case 2 had a common ancestor more than 14 years prior to the surgery. Case 4, the other suspected related isolate, was as distant from the surgical team member isolate as Case 2, with a 285 SNP differentiation.

Additionally, in order to determine epidemiologically related and unrelated isolates using WGST, a threshold was established from a set of related isolates for our study, using multiple samples taken from individual patients over time (Healthcare Control 9a and 9b, Case 3, 3a and 3b, Hospital Control 8a and 8b and Hospital Control 4a and 4b). The average pairwise distance between each set of related isolates was 2.3 SNPs with the exception of Case 3. Case 3, 3a and 3b were from the same patient over the span of one year and taken from multiple body sites. These three isolates contained 21 SNPs among them, with no more than 10 SNPs between any two isolates; however, this number is still within the predicted baseline mutation rate of *S. aureus*. Healthcare control samples 9a, 9b and 11 had two SNP differences separating them, indicating a likely recent transmission event, probably within the long-term care facility where both patients resided.

## Discussion

Our investigation determined that the surgical team member isolate was not the source of a transmission event and that no evidence of transmission occurred among the suspect cases. It is well established that humans are commonly colonized with *S. aureus*, with a fraction of the population being carriers of MRSA; it is estimated that 0.2-7% of inpatient hospital populations and upwards of 42.2% of hospital staff are MRSA carriers [28–30]. A recent study estimates *S. aureus* carriage among surgeons as high as 45.4% [31]. A high incidence of both methicillin-susceptible *S. aureus* (MSSA) and MRSA nasal carriage are exacerbating the increasing number of healthcare-associated *S. aureus* infections in patient care facilities [32]. Reducing patient colonization has been hypothesized to help in the prevention of MRSA transmission and infection [28, 29]. However, the impact of MRSA carriage by medical providers and hospital ancillary staff when compared to the MRSA nasal carriage rate of patients, in association with the incidence of MRSA nosocomial infections, is still not well understood but has been speculated as a source of patient infection[31, 33, 34]. Such provider to patient transmission would have significant liability, reimbursement and risk management implications for both providers and healthcare facilities. In this study, we used high-resolution techniques to compare a MRSA isolate carried by a surgical team member to the MRSA isolates of that surgical team member’s patients who had post-operative SSIs. WGST confirmed that transmission events did not occur between a healthcare provider and the surgical patients examined. In order to determine accurate epidemiological association, background control isolates from the hospital and community were compared to the suspected cluster isolates. By including these background isolates, if a transmission event had occurred, we would have observed a clear phylogenetic distinction between the background isolates and the isolates involved in the transmission event. However, we identified more than 17,000 parsimony informative SNPs among the 24 isolates and we did not observe a distinct grouping of the isolates that were originally thought to be surgery-associated transmission events.

WGST has been successfully used as a molecular epidemiology tool for numerous outbreak investigations [13–18, 35]. Our study further establishes the effectiveness of WGST for epidemiological investigations within a hospital setting, which is especially critical for rapid analysis during an ongoing outbreak. Interestingly, we identified a previously unrecognized transmission event outside the hospital that likely would not have been discovered without WGST. Furthermore, this study highlights the need for additional investigation to identify the source of hospital-acquired MRSA infections, including but not limited to medical providers, hospital staff members and the patient’s own nasal colonization, as well as the importance of source tracing to correctly identify an outbreak.

Implementing the reported mutation rate within our data disqualifies our hypothesis of a healthcare provider-related transmission, with at least 14 years of evolutionary time between the isolates from the surgical team member and the respective patients. Furthermore, the high resolution of WGST provides a genetic threshold to gauge relatedness between isolates collected from a single patient from multiple body sites and over time in comparison to isolates from unrelated patients. This genetic relatedness framework also allowed for the detection of a previously unrecognized MRSA transmission event. These data also suggest importation of strains circulating in other care facilities is a potential source for hospital-acquired infections.

A limitation of this study is the sequencing of a single colony from the suspected source. Recent research shows the bacterial diversity within a host can impede accurate depiction of transmission events [36]. Pathogen genetic diversity within a host can be widespread and may be due to numerous transmissions from multiple sources [36]. Furthermore, *S. aureus* studies reveal carriage of numerous sequence types within a single host likely due to multiple independent transmissions [37, 38]. In order to account for within-host diversity and reduce the risk of falsely interpreting results, it will be important to increase the sampling of isolates from the suspected source in future investigations. However, the fact that none of the three SSI isolates were related to one another provides evidence against a point source regardless of the single isolate sampling from the surgical team member.

## Conclusions

The lack of a MRSA transmission event between the colonized surgical team member and the surgical patients shown here has implications in the healthcare field, from infection prevention to hospital epidemiology and healthcare reimbursement. These findings were only apparent through the use of WGST. Low resolution strain typing methods like antibiogram profiles may lead to erroneous conclusions, and may instigate inappropriate, costly, and ineffective infection control measures as was the case here, as well as limit reimbursement for inpatient services from third-party payers (e.g., CMS) [39]. The accurate determination or repudiation of a healthcare-associated outbreak or cluster will prevent unnecessary, disruptive and costly infection prevention procedures and allow the implementation of these control measures when they are appropriate. With the increasing reality of clinical laboratory bacterial sequencing and accessible analysis tools, healthcare and public health systems would do well to maintain isolate and/or sequence repositories of local critical nosocomial pathogens to quickly detect and analyze possible hospital associated infection (HAI) outbreaks.

## Abbreviations

**MRSA:** Methicillin resistant *Staphylococcus aureus* **SSI:** Surgical site infection **SNP:** Single nucleotide polymorphism **WGST:** Whole genome SNP typing **WGS:** Whole genome sequencing **PCR:** Polymerase chain reaction **NASP:** Northern Arizona SNP pipeline **GATK:** Genome analysis toolkit **MLST:** Multi-locus sequence type **PFGE:** Pulse-field gel electrophoresis **ST:** sequence type **MSSA:** Methicillin resistant *Staphylococcus aureus* **HAI:** Hospital associated infection

## Author’s Contributions

CR analyzed the data and contributed to writing and revising the manuscript. KH contributed to the design of the study and sample collection. ED contribute to the design of the study, the analysis and writing and revising the manuscript. JB contributed to study design, processing samples and writing and revising the manuscript. JT contributed to writing the manuscript and the interpretation of the data. PK contributed to the study design, writing and revising the manuscript. DE contributed to the study design, writing and revising the manuscript. All authors read and approved the manuscript.

